# Agrin/Lrp4 signal constrains MuSK activity during neuromuscular synapse development in appendicular muscle

**DOI:** 10.1101/2021.05.07.443163

**Authors:** Lauren J Walker, Rebecca A Roque, Maria F Navarro, Michael Granato

## Abstract

The receptor tyrosine kinase MuSK, its co-receptor Lrp4 and the Agrin ligand constitute a signaling pathway critical in axial muscle for neuromuscular synapse development, yet whether this pathway functions similarly in appendicular muscle is unclear. Here, using the larval zebrafish pectoral fin, equivalent to tetrapod forelimbs, we show that like axial muscle, developing appendicular muscles develop aneural acetylcholine receptor (AChR) clusters prior to innervation. As motor axons arrive, neural AChR clusters form, eventually leading to functional synapses in a MuSK-dependent manner. Surprisingly, we find that loss of Agrin or Lrp4 function, which abolishes synaptic AChR clusters in axial muscle, results in enlarged presynaptic nerve endings and progressively expanding appendicular AChR clusters, mimicking the consequences of motoneuron ablation. Moreover, *musk* depletion in *lrp4* mutants partially restores synaptic AChR patterning. Combined, our results provide compelling evidence that, in contrast to axial muscle in which Agrin/Lrp4 stimulates MuSK activity, Agrin/Lrp4 signaling in appendicular muscle constrains MuSK activity to organize neuromuscular synapses. Thus, we reveal a previously unappreciated role for Agrin/Lrp4 signaling, thereby highlighting distinct differences between axial and appendicular synapse development.

## INTRODUCTION

Movement depends on coordinating development of functional synapses called neuromuscular junctions (NMJs) between motor axons and skeletal muscle. Prior to axon arrival, the central region of muscles exhibit a ‘prepattern’ of clustered acetylcholine receptors (AChR), which requires the receptor tyrosine kinase MuSK. Prepatterning is important for axon guidance, as axons navigate towards aneural AChR clusters on muscles and incorporate them into newly formed NMJs (Panzer et al., 2006). Disruption of prepatterning, such as in *musk* mutants, leads to exuberant motor axon outgrowth (Jing et al., 2009; Kim and Burden, 2008). Subsequently, as motor axons contact the muscle and transform into nerve endings, they release the glycoprotein Agrin which binds to the low-density lipoprotein receptor-related protein 4 (Lrp4) on the muscle membrane to stimulate phosphorylation of MuSK. Upon activation, MuSK initiates a downstream signaling cascade to cluster AChRs in apposition to axons, thereby forming neural synapses. While there are many additional proteins that modulate NMJ development, mutants of *lrp4*, *agrin*, and *musk* all fail to form NMJs in the mouse and zebrafish trunk, demonstrating their critical and conserved role in this process (Kim and Burden, 2008; Kim et al., 2008; Zhang et al., 2008; Jing et al., 2010; Remédio et al., 2016; Gribble et al., 2018). The muscular system is comprised of two divisions: axial and appendicular muscles. Axial muscles, such as the diaphragm, attach to the bones of the trunk whereas appendicular muscles move appendages, such as limbs. Extensive work on neuromuscular synapse development that identified the Agrin/Lrp4/MuSK pathway has focused predominantly on axial muscles. In contrast, significantly less is known about the cellular and molecular mechanisms critical for NMJ development in appendicular muscles.

Here, we employ the larval zebrafish pectoral fin, which is evolutionarily analogous to tetrapod forelimbs (Mercader, 2007), to study the process of neuromuscular synapse development in a paired appendage. In development, fin buds arise from mesenchymal protrusions oriented vertically just dorsal to the yolk and lateral to the second and third myotome (Grandel and Schulte-Merker, 1998). As the fin bud forms into the pectoral fin, it rotates approximately 90 degrees so the tip points caudally. Additionally, the fin migrates anteriorly, such that it is positioned posterior to the otic vesicle and anterior to the yolk by 60 hours post fertilization (hpf). At 120 hpf (5 days post fertilization (dpf)), pectoral fins are comprised of two antagonistic muscles, the abductor and adductor, separated by an endoskeletal disk (Fig. 1A,B). Each muscle consists of ~50-55 fast-twitch muscle fibers that extend longitudinally from the proximal fin base, where it attaches to the trunk, out to the distal tip of the fin. At the fin base the abductor and adductor muscles are each 2-3 muscle fibers thick, but muscles then thin out to be a single fiber layer throughout most of the fin (Thorsen et al., 2004). The abductor and adductor muscles are innervated by 4 distinct motor nerves, which we refer to here as nerves 1-4, with cell bodies in anterior spinal cord segments 3 through 6 (Myers, 1985; Thorsen and Hale, 2007). Motor axons enter the fin at a dorsal (nerves 1-3) or ventral (nerve 4) plexus to sort between the abductor or adductor muscles (Thorsen and Hale, 2007). Axons then progressively defasciculate as they grow towards the distal tip of the fin such that each muscle fiber is polyinnervated and motor axons create a patchwork pattern across the fin muscles. This innervation pattern remains unchanged until juvenile stages at three weeks (5.4-5.8 mm) when the muscles divide, nerves increase in arborization, and bone forms (Grandel and Schulte-Merker, 1998; Thorsen and Hale, 2007). The genetic-tractability, transgenic tools to label specific cell types, optical transparency suitable for live imaging, and behavioral readout of fin movement make the larval zebrafish pectoral fin an ideal vertebrate system to interrogate mechanisms of neuromuscular synapse development within appendicular muscles.

**Figure 1:**
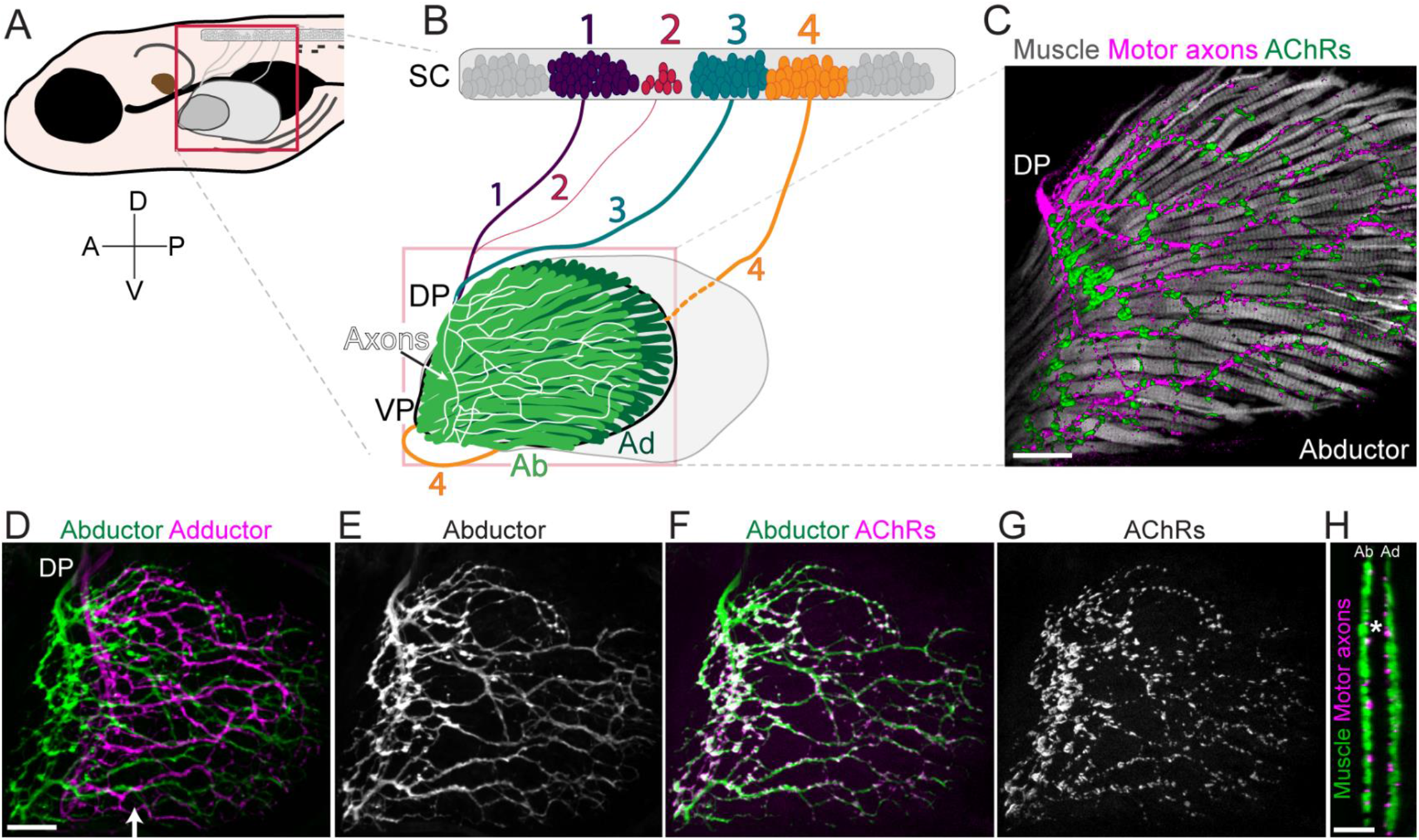
Pectoral fin anatomy. A) Schematic of 120 hour post fertilization (hpf; 5 days post fertilization) zebrafish larvae. Anterior (A) is to the left, dorsal (D) is up. B) Schematic of pectoral fin motor neuron innervation. Motor neuron cell bodies are in the anterior four segments of the spinal cord (SC). Nerves 1-3 enter the fin at the dorsal plexus (DP) while nerve 4 enters the fin at the ventral plexus (VP). All nerves innervate both the abductor (Ab) and adductor (Ad) muscles. C) Abductor innervation of 120 hpf *Tg(α-actin:GFP);Tg(Xla.Tubb:DsRed)* pectoral fin stained with α-bungarotoxin to visualize muscle fibers, axons, and acetylcholine receptors (AChRs), respectively. n = 7. D) Maximum projection of *Tg*(*mnx1*:*GFP*) axon innervation in pectoral fin. Abductor (green) and adductor (magenta) innervation patterns are pseudo-colored. E) Abductor innervation alone F) Abductor innervation with AChRs labeled with α-bungarotoxin G) AChR labeling alone. n > 77 wildtype pectoral fins for D-G. H) Cross-section of pectoral fin at approximate region marked by arrow in D. Asterisk marks endoskeletal disk that separates the two distinct muscles. Scale bars are 25 microns.

The axial trunk and the pectoral fin neuromuscular systems differ in several key ways. First, axon innervation of the axial muscles begins between 16-24 hours post fertilization (hpf) (Eisen et al., 1986) whereas in appendicular/fin muscles it is delayed by approximately 24 hours. Secondly, the trunk and the fin differ broadly in their muscle fiber anatomy. The trunk is comprised of several medial layers of fast-twitch (fast) muscle fibers and a single lateral slow-twitch (slow) muscle fiber layer that are arranged in repeating segments. In contrast, the fin muscles are only 1 fiber thick, comprised solely of fast fibers, and are approximately 2.5 times longer than trunk muscle fibers (Thorsen and Hale, 2007). Additionally, after exiting the spinal cord, axons in the trunk grow perpendicular to and along the center of muscle fibers. In contrast, the motor axons that innervate the pectoral fin grow through the body wall, sort at a plexus and then branch to create elaborate innervation patterns that can be perpendicular to and also parallel to muscle fibers in the fin. Axonal innervation of the fin is topographic, with axons from anterior spinal segments innervating the dorsal fin and axons from posterior segments innervating the ventral fin (Thorsen and Hale, 2007). Finally, in the trunk, aneural AChRs are present on slow muscle fibers, which are absent in the pectoral fin (Flanagan-Steet et al., 2005). Whether AChRs are prepatterned and how axon outgrowth occurs in the pectoral fin has not yet been described.

While many of the signals that mediate NMJ development have been well-characterized in axial muscles, whether the same cellular and molecular mechanisms underlie NMJ development within the complicated muscle arrangement of paired appendages has remained unclear. Here, we reveal that while some aspects of neuromuscular synapse development, such as prepatterning with aneural AChR clusters and the requirement for MuSK to form neural AChR clusters, are shared between the trunk and the pectoral fin, there are key muscle-specific differences in the fin. We show that both axonally released Agrin and Lrp4 on muscle cells are required to form the distributed patterning of AChR clusters in the fin. While *agrin* and *lrp4* mutants fail to form synapses in the trunk, in the pectoral fin they instead lead to the formation of giant AChR clusters and axonal innervation abnormalities. A developmental timecourse in *agrin* mutant fins reveals that these clusters likely arise from prepatterned AChR clusters that continue to grow over time. These giant clusters sequester navigating growth cones, thereby disrupting the innervation patterning. Partial depletion of *musk* moderately suppresses the formation of these clusters in *agrin* and *lrp4* mutants. Based on our results, we propose a model for NMJ formation in the appendicular fin muscle in which Agrin/Lrp4 signaling transitions MuSK from a prepatterning state to an axon-dependent, focal AChR clustering state. Without Agrin or Lrp4, MuSK remains active within prepatterned islands and AChR clusters grow. Importantly, this work exposes key differences in neuromuscular synapse development between axial muscles of the trunk and appendicular muscles of the pectoral fin.

## RESULTS

### Development of pectoral fin innervation is a tightly coordinated and dynamic process

At 120 hpf zebrafish pectoral fins have established a complex neuromuscular organization with motor axons forming an elaborate pattern across the abductor and adductor muscles. To visualize the organization between nerve and muscle within the fin we used transgenic *mnx1*:*GFP* or *Xla.tubb*:dsRed to label the majority if not all motor axons that innervate the pectoral fin and *α-actin:GFP* to label pectoral fin muscle fibers (Fig. 1A-C). Although the fin is comprised of both abductor and adductor muscles with independent innervation fields (Fig. 1D, H), for simplicity we include only the abductor innervation unless otherwise noted (Fig. 1E-G). Labeling of nicotinic AChRs with α-bungarotoxin (α-Btx) revealed hundreds of small, evenly spaced *en passant* neuromuscular synapses juxtaposed to motor axons (Fig. 1F-G). The largest postsynaptic AChR clusters are associated with the major nerve branch localized closest to the proximal fin base, while AChR clusters are smaller as the finer nerve branches defasciculate towards the distal tip of the fin musculature.

The developing pectoral fin is a dynamic structure. Previous work has described the development of the structure of the fin, from fin bud to adult (Yano et al., 2012; Siomava et al., 2018), as well as the innervation and musculature of the fin after 120 hpf (Thorsen and Hale, 2007), a timepoint at which the larval innervation pattern is relatively stable. Yet, to our knowledge the development of the zebrafish pectoral fin musculature and its complex innervation pattern prior to 120 hpf have not been described. To observe the process of muscle development and axonal innervation of the pectoral fin, we used long-term timelapse imaging of transgenic embryos to observe muscles (*Tg(α-actin:GFP*)) and axons (*Tg*(*Xla.tubb*:dsRed)) in developing zebrafish. By approximately prim-25 (36 hpf), motor axons from nerves 1-3 coalesce at what will form the dorsal plexus and nascent muscle fibers in the pectoral fin bud, located laterally to the axons, have just started expressing *α-actin*:*GFP* (Fig. 2A-B, Movie S1). Muscle fibers continue to divide and reorganize through the long-pec stage as the fin moves further medial, closer to the plane of the dorsal plexus, and motor axons begin to grow into the abductor and then adductor muscles beginning around the long-pec stage at 46 hpf. Concurrently, axons in nerve 4 make a sharp turn dorsally to innervate the fin via the ventral plexus. Thick axon bundles first grow perpendicular to muscle fibers near the proximal fin base, but subsequently axons turn posteriorly to grow mostly parallel to muscle fibers and towards the fin tip. As muscle fibers elongate, branching motor axons follow close behind to form a diffuse innervation network. By ~68 hpf, a simplified innervation pattern is established (Fig. 2B,G) that will become more complex through 120 hpf. Thus, the development of the innervation and musculature of the pectoral fin is a highly dynamic yet tightly coordinated process.

**Figure 2:**
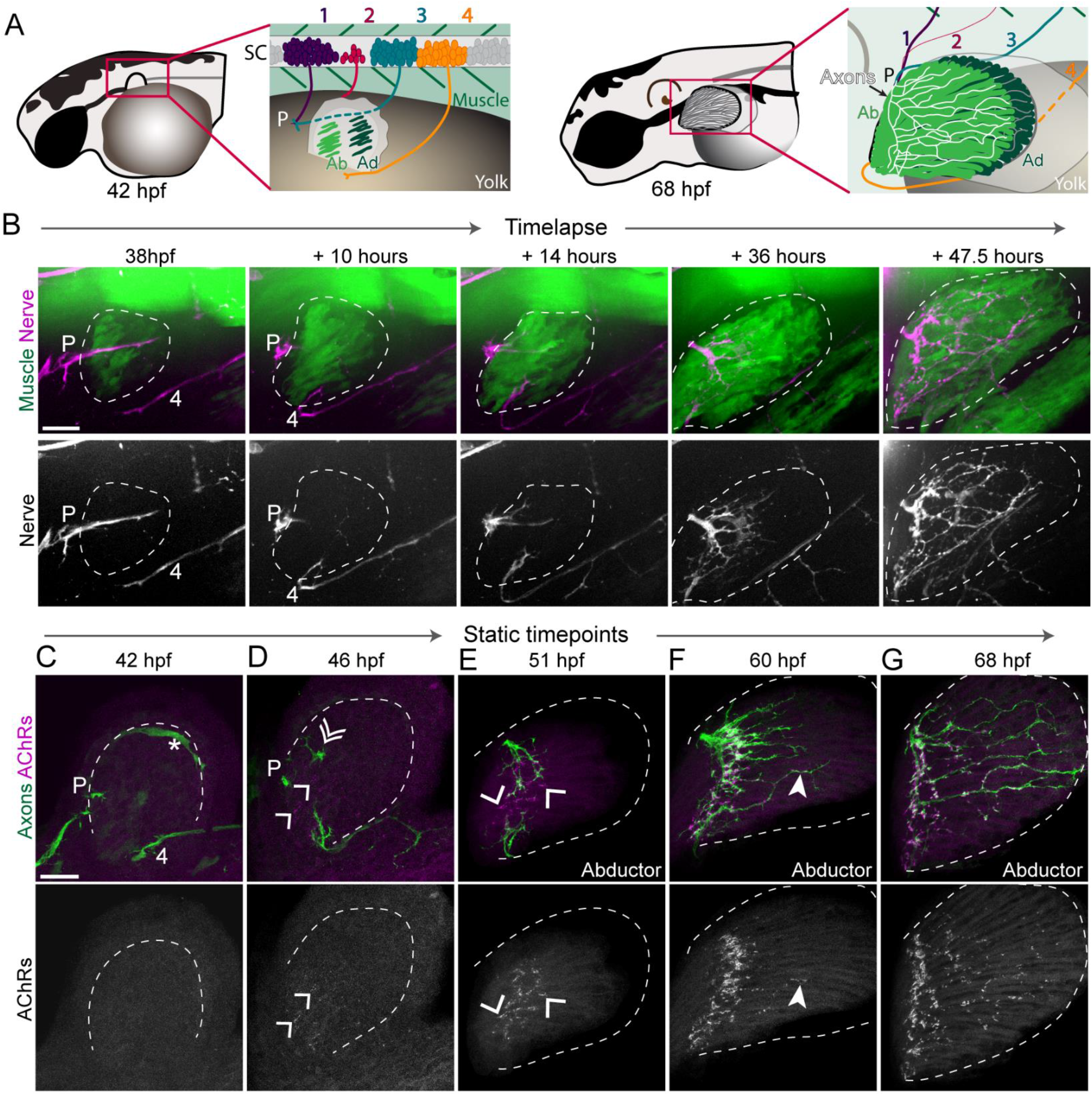
Development of pectoral fin innervation. A) Schematic of zebrafish larvae at 42 hours post fertilization (hpf), high pec stage, and 68 hpf, pec fin stage. Inset highlights motor neurons from spinal cord (SC) segments 1-4 and their corresponding axons projecting to the dorsal plexus (P) to innervate the abductor (Ab) and adductor (Ad) muscles of the pectoral fin. B) Maximum projection stills from timelapse imaging of *Tg(α-actin:GFP);Tg(Xla.Tubb:DsRed)* larvae to label muscles and axons, respectively. Dotted line outlines pectoral fin musculature. Axons converge at a dorsal plexus prior to innervating nascent muscle fibers. As muscle fibers elongate the axonal innervation pattern elaborates. n = 7 wild type animals. Static timepoints of *Tg*(*mnx1*:*GFP*) larvae stained with α-bungarotoxin to label acetylcholine receptors (AChRs). Nerve 4 (4) is labeled. C) At 42 hpf the pectoral fin bud is still lateral to the plexus, so axons and muscles are not yet in the same plane. Asterisk notes developing vasculature that is also labeled by *mnx1:GFP*. D) Axons have just grown past the plexus (adductor axons labeled in double arrowhead). Arrowheads point to aneural AChR clusters. E-G) Abductor innervation only. Axons occupy prepatterned clusters and induce new AChR clusters as they grow throughout the fin. Filled arrowhead points to fin axon branch already associated with AChR clusters. n = 5-10 per timepoint for C-G. Scale bars are 25 microns.

Pectoral fins start moving rhythmically as early as 3 dpf (Uemura et al., 2020), prompting us to examine when during development neuromuscular synapses in the pectoral fin form, and to what degree this process mirrors synapse development in axial muscle. To observe neuromuscular synapse development in pectoral fins we fixed transgenic *mnx1*:*GFP* larvae (to label presynaptic motor axons) at various timepoints and labeled AChRs with fluorescently-conjugated α-Btx. In vertebrates, ‘prepatterned’ AChR clusters form on muscle fibers prior to motor neuron innervation (Lin et al., 2001; Yang et al., 2001), but whether or when this occurs in developing appendicular muscle of pectoral fins has not been investigated. At the high-pec stage (~42 hpf), when the pectoral fin bud is located lateral to the nascent dorsal plexus and axons have not yet entered the fin, AChR clusters are undetectable (Fig. 2C). By 46 hpf, as axons have just started to grow past the plexus onto the fin musculature, we observe small AChR clusters near the base of the fin that are not yet associated with labeled axons (Fig. 2D). At 51 hpf, axons growing from the dorsal and ventral plexi extend towards each other along the fin base and innervate nearby AChR clusters, while remaining axon free or aneural AChR clusters at the not yet innervated medial region of the fin increase in size (Fig. 2E). By 60 hpf all AChR clusters at the proximal fin base are associated with axons (Fig. 2F).

The entry of axons into the fin initiates a second phase of neuromuscular synapse formation. As axons grow beyond the fin base and branch to extend along muscle fibers, new AChR clusters are formed. These new AChR clusters are always associated with axons (Fig. 2F-G). The number of AChR clusters increases as the innervation pattern becomes more complex, such that by 120 hpf axons are dotted with hundreds of regularly spaced AChR clusters that remain relatively small (<5 μm^2^) (Fig. 2G). In contrast, the clusters at the fin base that preceded axon outgrowth continue to increase in size, perhaps as they are innervated by additional axons, such that by 120 hpf the biggest AChR clusters in the fin (>20 μm^2^) are in this proximal region. Thus, neuromuscular synapse development in appendicular muscle of the pectoral fin mirrors that in axial muscle of the trunk, with a prepatterned phase and an axon-associated phase of synapse formation.

### MuSK signaling is required for neuromuscular synapse development in pectoral fin muscle

Given the similarities in AChR prepatterning and axon-associated AChR clustering between the trunk and the pectoral fin, we wondered if the well-established genetic pathway that regulates neuromuscular synapse development in axial muscle is also critical for this process in pectoral fin muscle. In vertebrate axial muscles, AChR prepatterning and the formation of neuromuscular synapses requires the receptor tyrosine kinase MuSK (DeChiara et al., 1996; Lefebvre et al., 2007). We first asked if *musk* is also required for AChR prepatterning within larval zebrafish appendicular muscle by staining sibling and *musk* mutant pectoral fins with α-Btx to label AChR clusters during the time of axon navigation into the fin. At the long-pec stage (46 hpf), when wildtype sibling animals have developed robust prepatterned aneural AChR clusters, staining of *musk* mutant pectoral fins failed to reveal any evidence of prepatterned AChR clusters (Fig. S1). To account for a possible delay in pectoral fin development or aneural cluster formation we also examined *musk* mutants at later timepoints. At 51 hpf and 60 hpf, the extent of axon growth in the fin is comparable between siblings and *musk* mutants. At these timepoints wild type axons have innervated prepatterned AChR clusters and new clusters have formed, while in *musk* mutant fins α-Btx staining remains diffuse and muscles lack discernible AChR clusters. Thus, as in trunk axial muscle, *musk* is required for AChR prepatterning in appendicular fin muscle.

Next, we examined the role of MuSK in the formation of neural AChR clusters characteristic of neuromuscular synapses. In the zebrafish trunk, motor axons branch and form synapses distributed along myofibers throughout the myotome. Consistent with previous work (Lefebvre et al., 2007; Jing et al., 2010), we find that at 120 hpf, trunk axial muscle fibers in *musk* mutants exhibit diffuse bungarotoxin staining with fewer and smaller AChR clusters, compared to sibling controls (Fig. 3 A-B, D). Similar to what we observe in trunk axial muscle fibers, appendicular muscle fibers in *musk* mutants display mostly diffuse bungarotoxin staining and lack wild type-like AChR clusters (Fig. 3C). While *musk* mutants exhibit some AChR clusters that form in apposition to axons, compared to sibling controls they are fewer and smaller in size. These circular clusters resemble the dystroglycan-dependent clusters that form on zebrafish axial muscle fibers in the absence of *musk* (Lefebvre et al., 2007). Moreover, compared to wild type controls, motor axons in *musk* mutant pectoral fins are less fasciculated, similar to what has been reported previously for axial muscle fibers in zebrafish and mouse lacking MuSK (Kim and Burden, 2008; Jing et al., 2009).

**Figure 3:**
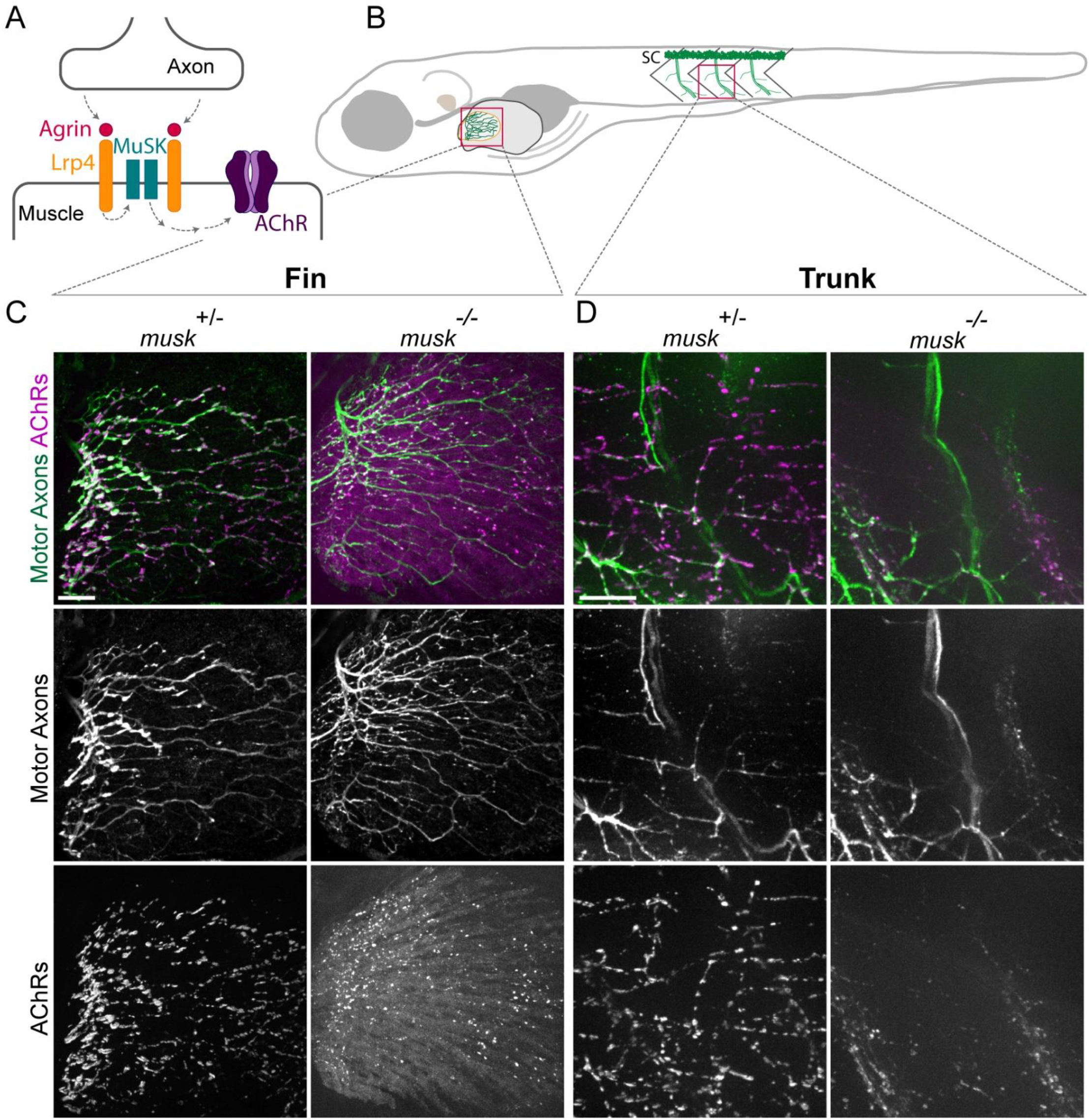
MuSK is required for pectoral fin neuromuscular synapse development. A) Schematic of Agrin/Lrp4/MuSK pathway B) Schematic of 120 hpf larval zebrafish. Red boxes outline regions of fin and trunk that were imaged for C-D. Spinal cord (SC). C) Abductor muscle innervation in the pectoral fin from 120 hpf larvae expressing *Tg*(*mnx1:GFP*) to label motor neurons and stained with α-bungarotoxin to label acetylcholine receptors (AChRs). *musk* heterozygous sibling pectoral fins exhibit an innervation pattern with numerous small AChR clusters (n = 45/45) while *musk* mutants have an exuberant innervation pattern with diffuse AChR signal throughout muscle fibers in the fin and some focal AChR clusters (n= 25/25). D) Trunk innervation from the same animals shown in C. *musk* mutants form fewer and smaller neuromuscular synapses. All images are maximum intensity projections that include the same number of slices for both genotypes. Scale bars are 25 microns.

Finally, we asked whether MuSK acts through its well-established downstream effector Rapsyn during appendicular neuromuscular development. Like *musk*, *rapsyn* is required for AChR clustering in mouse and zebrafish axial muscle (Gillespie et al., 1996; Ono et al., 2002). Similar to its role in axial muscle, *rapsyn* is required for AChR clustering in the pectoral fin, as *rapsyn* mutants display diffuse α-Btx signal throughout pectoral fin muscles (Fig. S2). Yet interestingly, in contrast to *musk* mutants, which have a defasciculated and overgrown pectoral fin axon patterning (Fig. 3C, 7B), we find that motor neuron innervation in the pectoral fin of *rapsyn* mutants is indistinguishable from wildtype (Fig. S2), similar to what has been observed in *rapsyn* mutant trunk innervation (Zhang et al., 2004; Gribble et al., 2018). Thus, *musk* and *rapsyn* are required for NMJ development in the pectoral fin, suggesting that postsynaptic mechanisms critical for synapse formation are shared between trunk and appendicular muscle.

### Axonal derived signals are critical for appendicular AChR patterning

Neuromuscular junction development requires precise, bidirectional coordination between axons and muscles. To determine the overall role of motor axon derived signals on the postsynaptic innervation pattern, we laser-ablated the motor neurons that innervate the dorsal portion of the pectoral fin muscle. Specifically, we ablated the cell bodies of motor neurons in spinal segments 3-6 at 42 hpf, prior to the growth of motor axons into the pectoral fin bud, and then re-ablated any newly formed motor neurons 24 hours later. Importantly, we left intact all motor neurons in spinal segment 4, which innervate the ventral muscle region of the fin, to serve as an internal control when comparing the innervation patterns of innervated and nerve-deprived muscle fibers. As expected, at 120 hpf AChR patterning in the innervated ventral region of the fin was indistinguishable compared to non-ablated controls, with small, evenly spaced AChR clusters in apposition to axons (Fig. 4A-C). Surprisingly, in the dorsal region of the motor neuron-ablated fin, which had never been innervated, fewer but much larger AChR clusters formed (Fig. 4C). These clusters were globular and evenly dispersed throughout the non-innervated musculature and differ vastly in size and distribution from those observed in the absence of MuSK or Rapsyn (Fig. 3; Fig. S2). Thus, while lack of the MuSK-dependent postsynaptic signal transduction machinery blocks the formation of AChR clusters almost completely, absence of axonal-derived signals results in unpatterned yet exuberantly sized AChR clusters, which going forward we refer to as giant AChR clusters. We conclude that axonal-derived signals are critical for AChR patterning and for limiting AChR cluster size.

**Figure 4:**
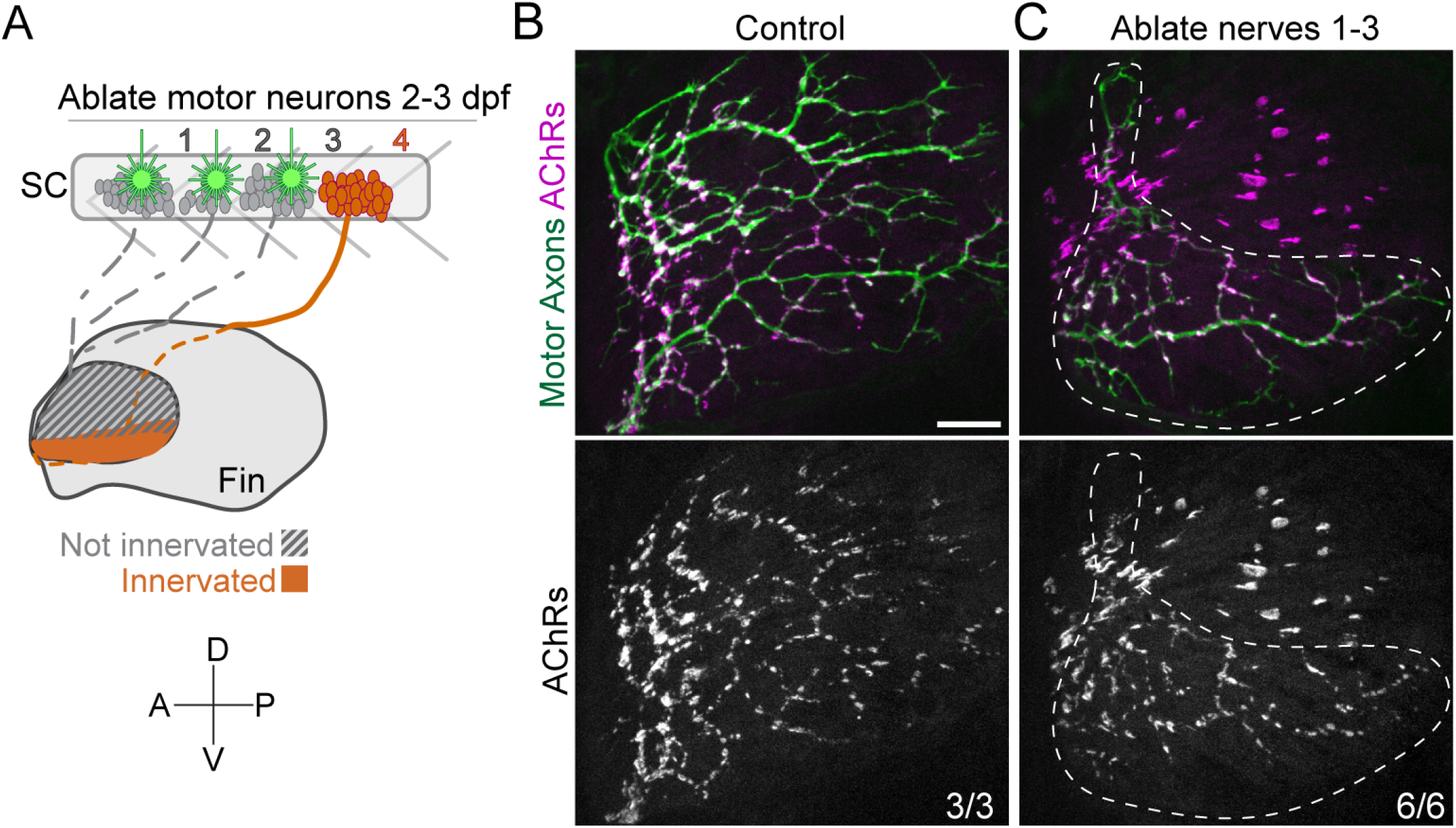
Pectoral fin muscles are predisposed to form large AChR clusters. A) Motor neuron cell bodies from spinal cord (SC) segments 1-3 were laser-ablated at 2 and 3 days post fertilization (dpf) to prevent motor axon innervation of the dorsal pectoral fin. A/P = anterior/posterior, D/V = dorsal/ventral B) Pectoral fins from control *Tg*(*mnx1*:*GFP*) larvae stained with α-bungarotoxin to label acetylcholine receptors (AChRs). C) Pectoral fin in *Tg*(*mnx1*:*GFP*) wild-type larvae after motor neuron ablation. The ventral fin was still innervated (outlined region) with input from unablated nerve 4. Non-innervated dorsal region of the fin has enlarged AChR clusters. N= 3/3 unablated controls, 6/6 wildtype pectoral fins with full or partial ablations at 120 hours post fertilization (5 dpf). Scale bar is 25 microns.

### Agrin and Lrp4 are required for appendicular neuromuscular development

The opposing consequences of blocking MuSK-dependent postsynaptic signaling versus eliminating all presynaptic signaling prompted us to examine the role of signaling components that activate MuSK. The heparan sulfate proteoglycan Agrin is an axon-derived signal that coordinates MuSK-dependent AChR clustering between nerve terminals and muscle fibers (Kim et al., 2008; Zhang et al., 2008), and zebrafish mutants lacking the motoneuron-derived Agrin isoforms lack synaptic AChR clusters along axial trunk muscle fibers (Fig. 5A) (Gribble et al., 2018). We therefore hypothesized that Agrin might play a similar critical role in inducing neural AChRs in pectoral fin muscle fibers. Throughout the fin of wild type siblings, hundreds of <5 μm^2^ AChR clusters are evenly distributed along appendicular muscles fibers. In contrast, AChR clusters in *agrin* mutants were reduced in numbers yet increased in size (>20 μm^2^) (Fig. 5B, E), indistinguishable from the giant AChR clusters we observed in nerve-deprived fins (Fig. 4C). To quantify the number of neural AChR clusters and their size distribution across genotypes, we focused on AChR clusters present on abductor muscle fibers, although quantification of adductor muscles across all genotypes revealed similar results (Fig. S3). Wildtype abductor muscle fibers exhibit 294.4± 21.5 α-Btx-positive AChR clusters per fin with a median cluster size of 3.9± 0.6 μm^2^ (Fig. 5C-E). In contrast, *agrin* mutant abductor muscle fibers have significantly fewer clusters per fin (51.6± 5.9, unpaired t-test p<0.0001) yet exhibit a vastly increased median cluster size of 18.4± 4.7 (t-test p<0.0001). Thus, in striking contrast to *agrin* mutant trunk muscle fibers that exhibit only small AChR clusters (median cluster size, siblings: 3.5 ± 0.7 μm^2^ (n=17), *agrin*^−/−^: 1.7 ± 0.7 μm^2^ (n=9), t-test p<0.0001), *agrin* mutant appendicular muscle fibers exhibit greatly enlarged AChR clusters.

**Figure 5:**
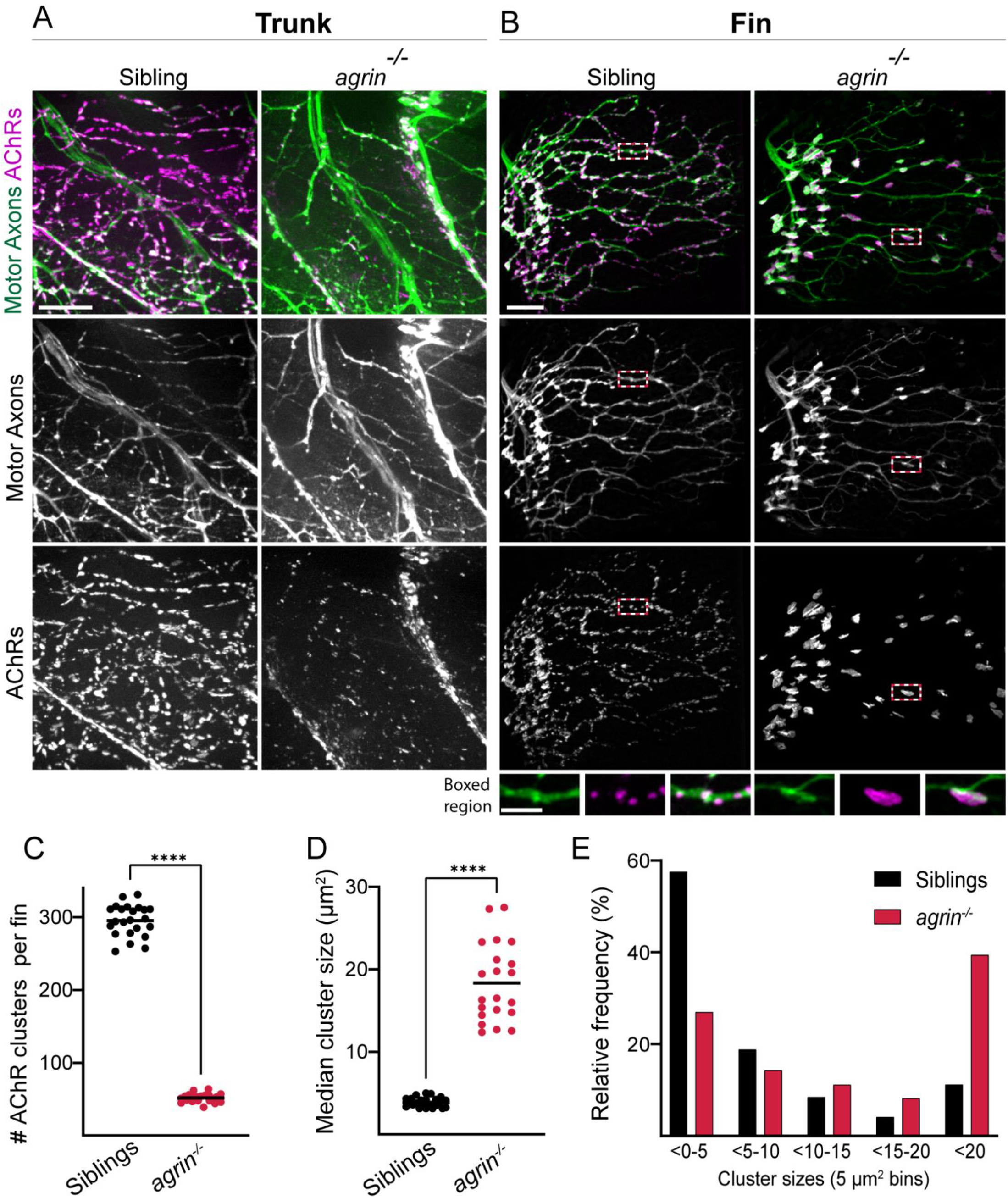
Agrin is required for correct axon innervation and AChR patterning in the pectoral fin. A) Trunk innervation in 120 hpf larvae expressing *Tg*(*mnx1:GFP*) to label motor axons and stained with α-bungarotoxin to label acetylcholine receptors (AChRs). Trunks in *agrin* mutants form fewer and smaller neuromuscular synapses. B) Abductor muscle innervation in the pectoral fin from the same animals shown in A. *Agrin* sibling animals exhibit an innervation pattern with numerous small AChR clusters while *agrin* mutants have swellings in the innervation pattern directly opposed to enlarged AChR clusters throughout muscle fibers in the fin. Scale bar is 25 microns. Inset from boxed region shows even distribution of small magenta AChR clusters in siblings while mutants have large AChR clusters that colocalize with green axon swellings. Inset scale bar is 10 microns. All images are maximum intensity projections that include the same number of slices for both genotypes. Quantification of the number of AChR clusters per fin (C) and the median cluster size per fin (D). E) Histogram of the distribution of AChR cluster sizes across all fins quantified (5 square micron bins). ****; P<0.0001, t-test. n = 23 (siblings), 21 (mutants).

The strikingly divergent appendicular NMJ phenotypes observed in *musk* mutants that display an almost complete loss of AChR clusters versus the giant AChR clusters present in *agrin* mutants prompted us to examine the role of the Agrin receptor Lrp4. Upon binding Agrin, Lrp4 induces MuSK phosphorylation to initiate synaptic differentiation (Kim et al., 2008; Zong et al., 2012). To determine whether *lrp4* mutants display the phenotype exhibited by its canonical ligand Agrin or by its co-receptor MuSK, we examined the role of Lrp4 in appendicular neuromuscular development. Like *musk* and *agrin*, *lrp4* is required for neuromuscular junction development in axial muscles of the zebrafish trunk, as mutants form fewer and smaller synapses (Remédio et al., 2016). Identical to the *agrin* mutant fin muscle phenotype, and in contrast to the *lrp4* mutant phenotype in axial trunk muscles (Remédio et al., 2016), at 120 hpf *lrp4* mutant pectoral fin muscles displayed large AChR clusters (Fig. 6A-B, D-F). Moreover, in fins of *lrp4* mutants lacking the intracellular domain (Saint-Amant et al., 2008) we also observe giant AChR clusters (Fig. S4), providing compelling evidence that Lrp4 acts through a ligand-dependent mechanism to regulate appendicular NMJ development. Despite the abnormal AChR clusters in *lrp4* mutants, we failed to detect differences in the morphology of muscles or muscle fibers in pectoral fin muscles labeled with *Tg(α-actin:GFP)* indicating that in these mutants appendicular muscle development is unaffected (Fig. S4). Interestingly, co-labeling of AChR clusters and muscle fibers reveals that the enlarged clusters in *lrp4* mutants often nestle between adjacent muscle fibers but can also span across multiple fibers, similar to the enlarged clusters near the proximal fin base in wild type pectoral fins, suggesting extrinsic coordination between muscle fibers to form or stabilize these giant AChR clusters. Finally, expressing Lrp4 using a muscle specific promoter *Tg(α-actin:lrp4-GFP*) (Gribble et al., 2018) in otherwise *lrp4* mutant animals fully restored neuromuscular synapse development in appendicular muscle fibers (Fig. 6C-F), indicating that Lrp4 functions in muscle. Thus, loss of Agrin or Lrp4, while associated with a significant reduction in neural AChR cluster numbers in both axial and appendicular muscle, also leads to an increase in AChR cluster size on appendicular muscle, distinct from their mutant phenotypes in axial muscle.

**Figure 6:**
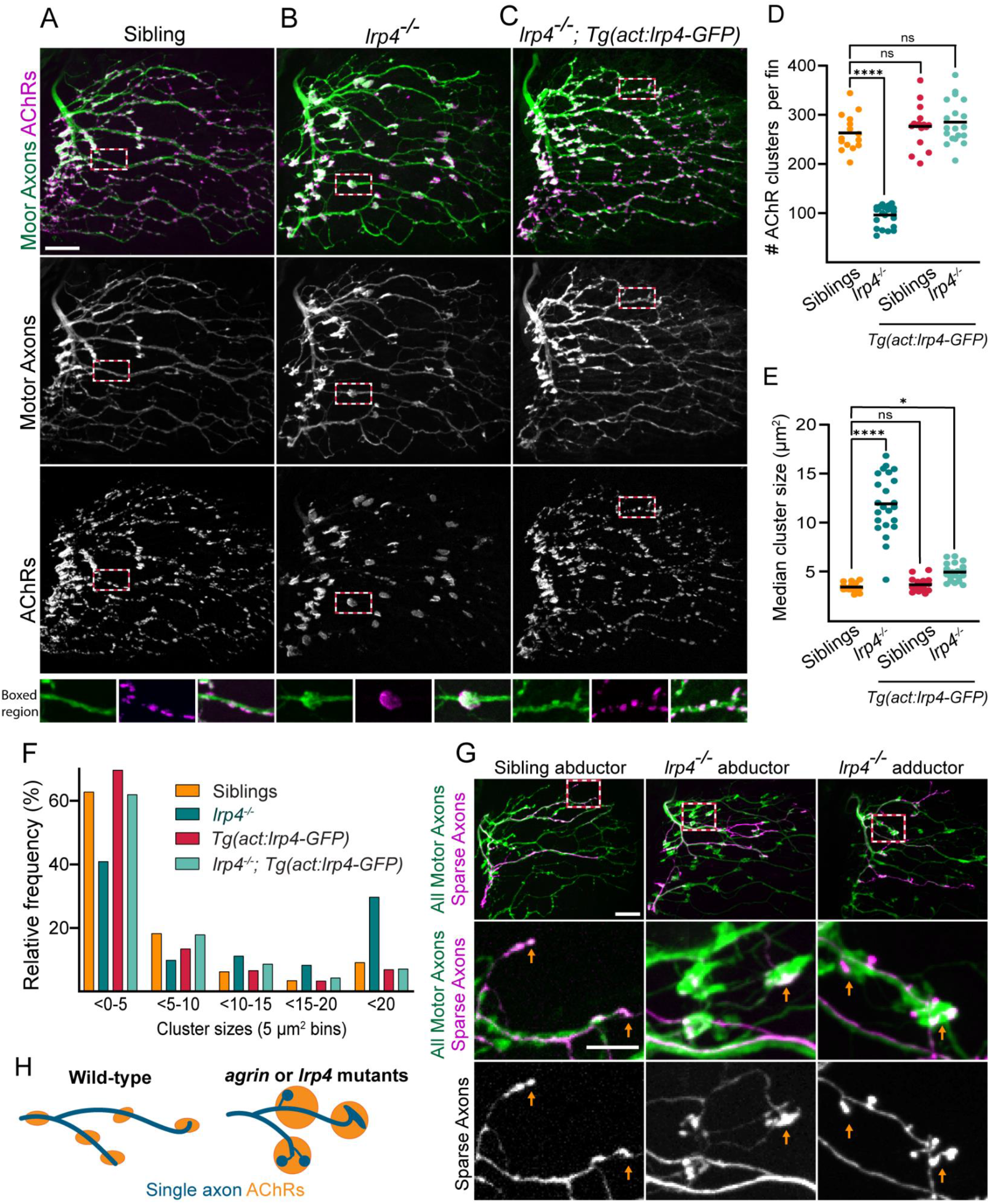
Lrp4 is required for correct axon innervation and AChR patterning in the pectoral fin. Abductor muscle innervation in the pectoral fin in 120 hours post fertilization larvae expressing *Tg*(*mnx1:GFP*) to label motor neurons and stained with α-bungarotoxin to label acetylcholine receptors (AChRs). A) *lrp4* sibling animals exhibit innervation patterns with numerous small AChR clusters. B) *lrp4* mutants exhibit abnormal swellings in the innervation pattern directly opposed to large AChR clusters. C) Expression of *lrp4*-*GFP* in muscles of *lrp4* mutants is sufficient to rescue the mutant innervation pattern. Scale bar is 25 microns. Dotted boxes outline region shown in insets (scale bar of insets is 10 microns). Quantification of the number of AChR clusters per fin (D) and the median cluster size per fin (E). F) Histogram of the distribution of AChR cluster sizes across all animals quantified (5 square micron bins). n = 15 (siblings), 23 (*lrp4* mutants), 16 (siblings plus *Tg(act:lrp4-GFP)*, 20 (*lrp4* mutants plus *Tg(act:lrp4-GFP)*. G) Sparse labeling of axons injected with *mnx1*:*mKate*, with all motor axons labeled with *Tg*(*mnx1:GFP*). Sparse labeling does not label entire ‘swelling’ but axon ends appear bulbous, indicating that multiple axons contribute to the abnormal innervation swellings in *lrp4* mutants. Orange arrowheads point to axon endings. Dotted box outlines insets. n = 11/12 (siblings), 7/7 (mutants) Scale bar is 25 microns. H) Schematic summarizing axonal organization and AChR clusters. ns= not significant, *p<0.05, ****p<0.0001, one way ANOVA with Dunnett’s multiple comparisons test.

### Agrin/Lrp4 regulates the size and patterning of appendicular neuromuscular synapses

The giant AChR clusters we observe on appendicular muscle fibers of nerve-deprived pectoral fins as well as in *agrin* and *lrp4* mutants resemble what has previously been described in *Xenopus* and chick muscle cells grown in the absence of axons as AChR ‘hot spots’, which disperse upon innervation (Bekoff and Betz, 1976; Moody-Corbett and Cohen, 1982; Peng, 1986). We therefore asked whether these giant AChR clusters we observe in *agrin* or *lrp4* mutant pectoral fins are caused by the lack of axonal innervation. Analysis of the axonal innervation pattern in wildtype and mutant animals at 120 hpf revealed that in both *agrin* and *lrp4* mutants the overall innervation pattern and extent of axon growth in pectoral fins is comparable to that of wild type, indicating that overall axon guidance mechanisms are largely intact. However, unlike in wild type siblings, in both *agrin* and *lrp4* mutant fins we observe large axonal ‘swellings’ (Fig. 5B; 6B). These axonal swellings co-localize with the enlarged postsynaptic AChR clusters, supporting the notion that, rather than being aneural AChR hot spots, these giant AChR clusters are indeed innervated and do not disperse upon axon contact. Moreover, expressing Lrp4 using a muscle specific promoter *Tg(α-actin:lrp4-GFP*) (Gribble et al., 2018) in otherwise *lrp4* mutant animals fully suppressed formation of presynaptic axonal swellings (Fig. 6C-F), indicating that muscle-derived Lrp4 signaling plays a role in establishing both presynaptic and postsynaptic neuromuscular synapse patterning.

To further investigate the nature of these axonal swellings, we used the Znp-1 antibody against the presynaptic marker Synaptotagmin 2. In sibling control pectoral fins, the Znp-1 signal concentrates at α-Btx-positive postsynaptic areas, demarcating presumptive presynaptic sites. Similarly, in *agrin* mutant pectoral fins, the Znp-1 signal colocalizes with the giant AChR clusters (Fig. S5). Likewise, we assessed the localization of the postsynaptic protein Dystrophin (Dmd) by analyzing a gene trap line (*dmd-citrine)* that labels endogenous Dmd (Ruf-Zamojski et al., 2015). The Dmd-citrine signal in sibling control pectoral fins is diffuse, with concentrations between muscle fibers and at synaptic regions, which we confirmed by co-labeling with fluorescent α-Btx. Similarly, in *lrp4* mutant pectoral fins, the Dmd-citrine signal is concentrated at regions marked by presynaptic axonal swellings and enlarged postsynaptic AChR clusters (Fig. S6). Thus, while in *agrin* and *lrp4* mutants AChR cluster size and distribution is altered, both presynaptic and postsynaptic proteins are recruited to these giant clusters suggesting that they are *bona fide* synapses.

We next examined the prominent presynaptic ‘swellings’ in *lrp4* and *agrin* mutants that form in opposition to enlarged AChR clusters. The *mnx1*:*GFP* or *Xla.tubb*:*dsRed* transgenic lines both label the entire population of motor axons in the pectoral fin, precluding us from visualizing the nature of these presynaptic swellings at the single axon level. These innervation swellings could be formed by 1) local distension or an increase in diameter of individual axons, 2) many individual axons forming synapses in a single spot on a muscle fiber, 3) abnormally enlarged axonal endings in a single spot on a muscle fiber, or 4) individual axons looping continuously in a single spot on a muscle fiber. To determine how individual axons contribute to the innervation swellings, we employed a sparse labeling strategy using *mnx1:m*Kate to visualize individual axons in the context of the entire population of motor axons (*mnx1:GFP*). We screened for larvae that expressed *mnx1*:*mKate* in only a few of the motor neurons that innervate the pectoral fin. In wild type siblings, we find that single axons branch and fasciculate with other axons to form complex patterns. Axons terminate abruptly, with endings that are approximately the same diameter as the rest of the labeled axon (Fig. 6G-H). In *lrp4* mutants, most individually labeled axons exhibit similarly complex trajectories, are similar in diameter to sibling controls, and occasionally form simplified endings. However, we also observed individual *mnx1:Kate*-positive axons that form bulbous and swollen structures. These globular endings of *mnx1*:*mKate* axons were part of larger *mnx1:GFP* swellings, indicating that many independent axons contribute to these swellings. As these swellings co-localize with α-Btx (Fig. 6B), we conclude that despite their abnormal morphology, they represent presynaptic terminals. This result strongly suggests that during appendicular neuromuscular development Agrin/Lrp4-dependent signaling not only promotes the formation postsynaptic AChR clusters but also limits their size, possibly by dispersing nascent AChR clusters or limiting cluster growth. In addition, Agrin/Lrp4 signaling also influences presynaptic patterning. Independent of the precise mechanisms by which Agrin and Lrp4 regulate pre and postsynaptic development, our data reveal that the role of Agrin and Lrp4 in zebrafish appendicular fin is distinctly different from its well-characterized function in trunk axial muscle.

### *musk* depletion partially suppresses the *lrp4* giant AChR cluster phenotype

Our results reveal that, unlike in axial trunk muscle of mice and zebrafish in which *musk*, *lrp4*, and *agrin* mutants all phenocopy each other, in appendicular muscle of the zebrafish pectoral fin loss of *agrin* or *lrp4* results in a phenotype that is distinct from *musk* mutants. Specifically, in *musk* mutants, appendicular muscle displays an almost complete loss of AChR clusters, while in *agrin* and *lrp4* mutant appendicular muscle, albeit exhibiting reduced numbers of AChRs, display a prominent, strikingly divergent phenotype characterized by enlarged AChR clusters. Combined, this led us to first ask whether, in the context of appendicular NMJ development, Agrin and Lrp4 act through MuSK, similar to what is observed in axial trunk muscle. Indeed, we find that *musk*;*lrp4* double mutants recapitulate the *musk* mutant phenotype as they fail to cluster AChRs and display axon overgrowth, confirming that Lrp4 acts through MuSK in NMJ development in both axial and appendicular muscle (Fig. 7A-B).

**Figure 7:**
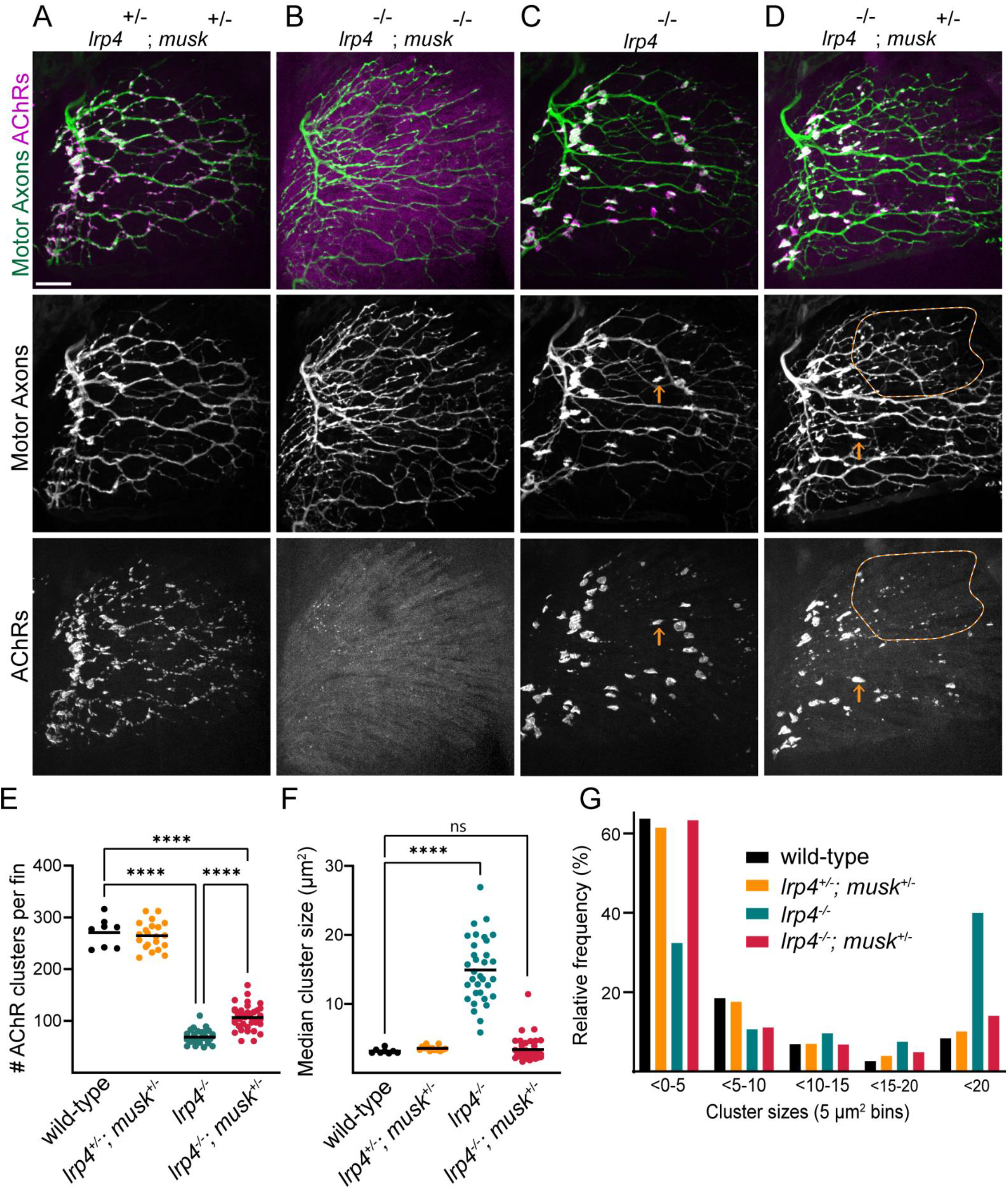
Musk depletion partially suppresses *lrp4* mutant phenotype. A) *lrp4^+/−^;musk^+/−^* trans heterozygotes have an innervation pattern indistinguishable from wild type while B) *lrp4;musk* double mutants phenocopy *musk* mutants with defasciculated axonal patterning labeled with Tg(*mnx1*:*GFP*) and diffuse acetylcholine receptor (AChR) staining as labeled by bungarotoxin. C) *lrp4* mutant motor axons have swellings in their innervation pattern that are opposed to large AChR clusters. D) While *lrp4* mutants that are heterozygous for *musk* (*lrp4^−/−^;musk^+/−^*) still have some large AChR clusters similar to *lrp4* mutants (orange arrows), they also have regions of the fin with smaller AChR clusters (orange dotted region). Quantification of E) the number of AChR clusters per fin, F) the median cluster size per fin (square microns), and G) the distribution of cluster sizes (5 square micron bins, ****p< 0.001, one-way Anova with Sidak’s (E) or Dunnett’s (F) multiple comparisons test. n= 8 (wild type), 21 (*lrp4^+/−^;musk^+/−^*), 4 (*lrp4^−/−^;musk^−/−^*), 33 (*lrp4*^−/−^), 39 (*lrp4^−/−^;musk^+/−^*).

As MuSK is necessary for both aneural and neural AChR clustering in the pectoral fin in both wild type and *lrp4* mutants and MuSK expression is sufficient to induce AChR clusters (Kim and Burden, 2008), we hypothesized that MuSK drives the formation of the enlarged AChR clusters in the absence of *agrin* or *lrp4* mutants. This would suggest, unexpectedly, that in appendicular muscle Agrin/Lrp4 may restrict MuSK function. If so, we would predict that dampening MuSK activity in *lrp4* mutants would suppress the giant AChR cluster phenotype. To this end, we examined *lrp4* mutants that lack one copy of *musk* (*lrp4^−/−^;musk*^−/+^). Indeed, *lrp4* mutant animals that are also heterozygous for *musk* exhibit a less severe phenotype than *lrp4* homozygous mutants. While fins in *lrp4^−/−^;*musk^+/−^ larvae still contained some giant AChR clusters, portions of the fins in these animals also contained smaller, evenly-dispersed neural clusters that resemble the sibling patterning. Additionally, the portions of the fin with smaller AChR clusters also lacked presynaptic axonal swellings. When compared to *lrp4^−/−^* mutants, *lrp4^−/−^;*musk^+/−^ mutants displayed an increase in the number of α-Btx-positive AChR clusters per fin (Fig. 7E), a rescue of the median cluster size (Fig. 7F), and a rescue of the overall distribution of cluster sizes in the fin (Fig. 7G). In *lrp4^−/−^;*musk^+/−^ pectoral fins, the giant clusters that formed often were closer to the proximal fin base, similar to the earlier-formed prepatterned clusters (Fig. 7D). In contrast, smaller clusters were often found in the distal fin, where AChR clusters formed later. This further supports the idea that Agrin/Lrp4 signaling restrains MuSK activity within appendicular muscle.

### Agrin/Lrp4 signaling restrains MuSK activity during appendicular NMJ development

To further explore the idea that, unlike in axial muscle, in appendicular NMJ development Agrin plays a critical role in restraining MuSK activity, we examined the progression of appendicular NMJ development in siblings and *agrin* mutants. We hypothesized that prior to the arrival of motor axons, the Agrin-independent formation of aneural prepatterned AChR clusters should be indistinguishable between wild type and *agrin* mutants. Indeed, we find that at 46 hpf (the long-pec stage), while axons are sorting at the dorsal plexus prior to growing into the fin, *agrin* mutant fins are prepatterned with AChR clusters and are indistinguishable from siblings (Fig. 8A). Moreover, like sibling controls, navigating axons in *agrin* mutants tend to occupy the prepatterned region near the proximal fin prior to extending towards the distal fin (Fig. 8B). Subsequently, in wild type, the arrival of motor axons and the release of Agrin induces the formation of small neural clusters, akin to the process previously described in axial muscle (Panzer et al 2006).

**Figure 8:**
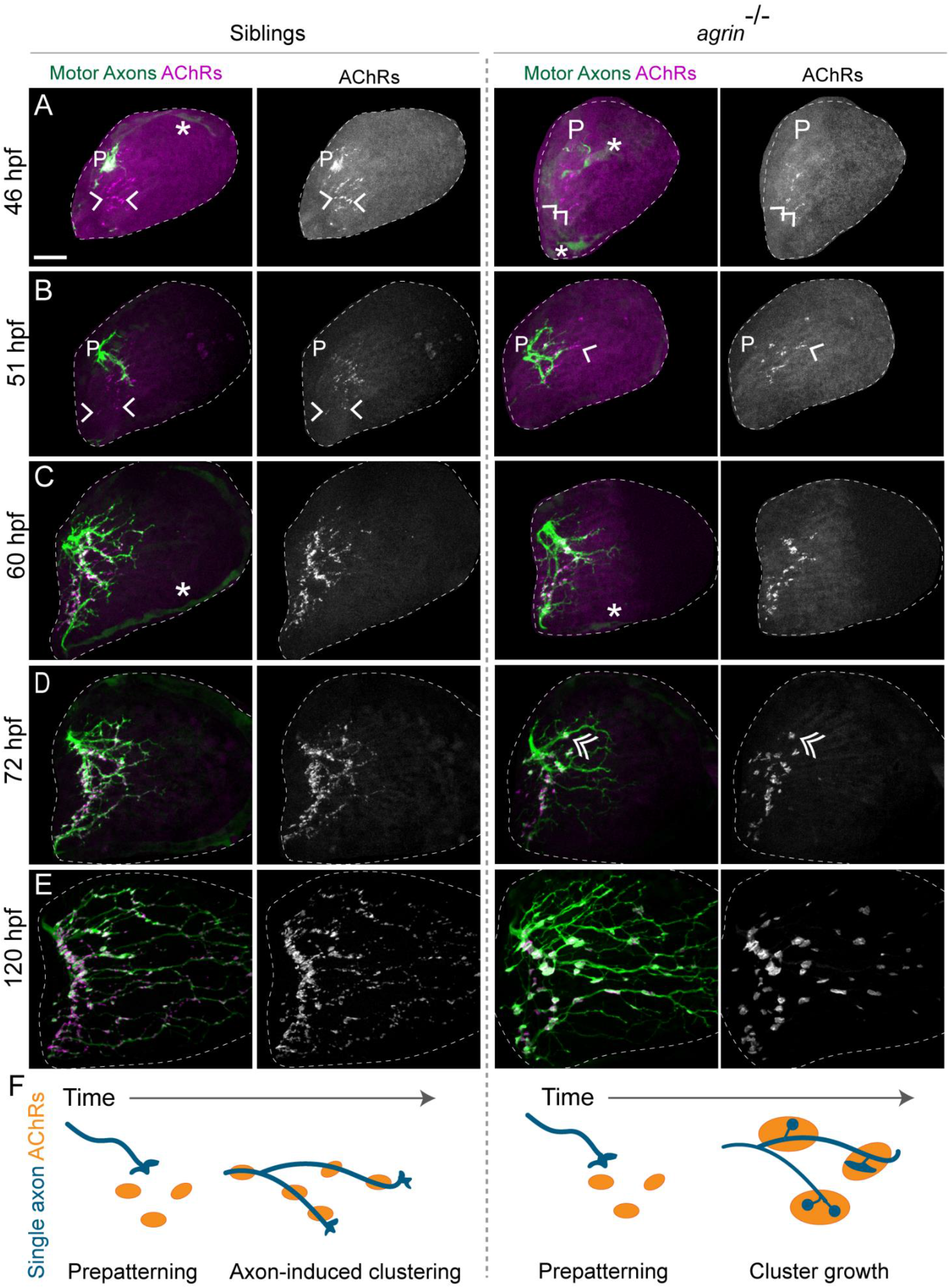
Agrin restricts presynaptic terminal and neural AChR cluster size. Developmental timecourse from *Tg*(*mnx1*:*GFP*) larvae to label motor neurons stained with α-bungarotoxin to label acetylcholine receptors (AChR). A-B). At 46 and 51 hpf, axons growing from the dorsal plexus (P) have not yet innervated all prepatterned AChR clusters (arrowheads). Asterisks note endothelial or endoskeletal cells labeled in the green channel. C-E) While small clusters are added as axons grow into the pectoral fin in sibling animals, clusters mainly increase in size in *agrin* mutants. D) Double arrow points to presynaptic swelling colocalized with an AChR cluster. Only abductor innervation is shown, with fin area outlined in dotted white line. Scale bar is 25 microns. G) Schematic summarizing developmental timecourse. Both siblings and *agrin* mutants look similar during prepatterning stage. Incoming axons induce small AChR clusters in sibling animals while in *agrin* mutants AChR clusters and axonal swellings increase in size over time. n= 4-10 animals per genotype per timepoint.

If nerve derived Agrin indeed restrains MuSK activity, we predicted that in *agrin* mutants these initially aneural clusters would retain their size (or even grow), despite the arrival of motor axons. We also predicted that lack of Agrin-mediated local MuSK activation would result in the failure to form new small neural clusters that normally emerge along growing axons. Indeed, at 60hpf (pec fin stage) the difference in cluster size, number, and distribution between genotypes is prominent. In wild type siblings, the AChR cluster field mirrors the innervation pattern. As motor axons continue to grow further towards the distal fin, even the furthest-reaching navigating axons are associated with small AChR clusters, suggesting that as axons grow, new clusters are rapidly formed. In contrast, in *agrin* mutants, AChR clusters have increased in size and remain globular (Fig. 8C). Unlike in wild type siblings, in *agrin* mutants presynaptic swellings apposed to AChR clusters become apparent by 72 hpf, with stretches of axon deprived of any discernible AChR clusters (Fig. 8D). This observation supports the idea that navigating growth cones are inappropriately attracted to and sequestered by these prepatterned ‘islands’. Between 72 and 120 hpf, both sibling and mutant axons continue to grow to occupy the entire muscle territory. In wild type siblings, the majority of the AChR clusters remain small (below 5 μm^2^) and evenly-dispersed throughout the appendicular muscle. In contrast, in *agrin* mutants both presynaptic axonal swellings and neural AChR clusters continued to grow throughout the entire appendicular muscle (Fig. 8D-F). Thus, lack of *agrin* leads to a progressive size increase of synaptic AChR clusters, consistent with the idea that in wild type appendicular muscle Agrin functions to counteract or to restrain MuSK activity. Together, our results suggest a model for appendicular NMJ development in which Agrin/Lrp4 signaling, different from its sole role in axial muscle to activate MuSK and promote NMJ development, also restrains MuSK activity to properly size and pattern neuromuscular synapses.

## DISCUSSION

Axial muscles of the mouse diaphragm or the larval zebrafish trunk have been major *in vivo* models to study neuromuscular synapse development. Genetic studies using these models have converged on an evolutionarily conserved, canonical pathway in which the axonally released glycoprotein Agrin binds its receptor Lrp4 on the muscle membrane to locally activate the receptor tyrosine kinase MuSK and cluster AChRs. Here, we employ the larval zebrafish pectoral fin as a genetically-tractable model system in which to study the development of the neuromuscular system within appendicular muscle. We use static and live-imaging to describe the coordinated growth of nascent muscle fibers, motor axon patterning, and postsynaptic AChR clustering in the fin. Using this framework, we provide compelling *in vivo* evidence that Agrin and Lrp4 play an additional, previously unappreciated role to regulate neuromuscular synapse development in appendicular muscle. We report that Agrin, Lrp4, and MuSK are required for the formation of neural AChR clusters in appendicular muscle, similar to their roles in axial muscle. In addition, we find that Agrin and Lrp4 play a second role to suppress the growth of aneural AChR clusters selectively in appendicular muscle. Consequently, only *lrp4* and *agrin* but not *musk* mutants form abnormally large postsynaptic AChR clusters in the pectoral fin. In addition, Agrin/Lrp4 signaling influences presynaptic axonal patterning, as in *lrp4* and *agrin* mutants multiple axons inappropriately innervate these giant AChR clusters. The formation of these abnormal synapses can be partially suppressed by depleting *musk*, providing compelling genetic evidence that Agrin and Lrp4 constrain MuSK activity to establish appropriate neural synapse formation in appendicular muscle. Thus, our work reveals a key difference in the regulation of neuromuscular synapse development between two major divisions of the muscular system, i.e. axial and appendicular muscles.

### Key differences between axial versus and appendicular NMJ development

In contrast to the neuromuscular system in axial muscle, little is known about the steps by which the complicated innervation pattern of appendicular muscle found in paired appendices including the pectoral fin arises, the genetic pathways that control it, nor how it might compare to NMJ development in axial muscles. Our work reveals key differences in neuromuscular synapse development between axial muscle in the zebrafish trunk and appendicular muscle in the pectoral fin. First, innervation of the trunk myotome occurs much earlier than the pectoral fin; trunk motor axon outgrowth begins at 16hpf (Panzer et al., 2006), whereas we first observe axons sorting at the fin plexus around 46 hpf. Secondly, both the trunk and the pectoral fin musculature are prepatterned with MuSK-dependent aneural AChR clusters. Somewhat surprisingly, prepatterned AChR clusters only form on adaxial, slow muscle fibers in the trunk (Flanagan-Steet et al., 2005; Panzer et al., 2006) and have not been detected in fast muscle fibers of the trunk, while in the pectoral fin, which lacks slow muscle fibers, prepatterned AChR clusters form exclusively on fast muscle fibers. This demonstrates that both slow and fast muscle fibers have the capacity to develop AChR prepatterning, thus it will be interesting to determine the mechanisms that selectively promote AChR prepatterning in trunk slow muscle and/or suppress it in adjacent fast muscle fibers. Third, in both zebrafish trunk and mouse diaphragm muscle, prepatterned AChRs are restricted to the central region of muscle fibers. We find that AChR prepatterning in pectoral fins is also restricted, but to the proximal region of individual muscle fibers. In the absence of live cell imaging, we cannot exclude the possibility that prepatterned AChR clusters initially arise in the ‘center’ of nascent pectoral fin muscle fibers, and, as these fibers elongate, clusters rapidly become off-center and shift towards the proximal region of muscle fibers. Independent of this possibility, our analysis of appendicular NMJ development reveals that AChR prepatterning is not strictly restricted to the center but instead can localize to other muscle fiber areas.

Finally, like AChR prepatterning, early axial motor axon outgrowth is confined to the center of the myotome and navigating growth cones contact prepatterned AChR clusters that form in this region. Similar to the trunk (Panzer et al., 2006), our developmental timepoints suggest that early extending axons selectively grow towards prepatterned muscle regions. Yet beyond the prepatterned region, axons in the pectoral fin form intricate patterns across muscle fibers. This growth pattern is more similar to later axon outgrowth in the trunk in which motor axons branch to innervate fast muscle fibers deep in the myotome (Beattie, 2000). These differences in prepatterning and axon outgrowth may be a consequence of the anatomy of the pectoral fin, in which axons converge at a dorsal or ventral plexus prior to topographically innervating longitudinal muscle fibers. Indeed, pectoral fin neuroanatomy is similar to that of the muscles of tetrapod forelimbs, in which axons sort at the brachial plexus to innervate distinct muscles. While we identify a conserved requirement for MuSK to establish prepatterning across muscles, these key anatomical and developmental differences between appendicular and axial muscles indicate that the signals that determine the location of AChR prepatterning and direct axon pathfinding are differentially regulated in appendicular muscle, leading to open questions regarding additional molecular mechanisms and pathway components that orchestrate NMJ development selectively in appendicular muscle.

### A dual role for Agrin/Lrp4 signaling in appendicular neuromuscular synapse development

Consistent with previous work, we find that loss of Agrin and Lrp4 leads to a significant reduction of neural AChR clusters on appendicular muscle fibers, demonstrating a critical and conserved role for both genes in promoting the formation these synaptic AChR clusters (Fig. 5C, Fig. 6D). Examining *agrin*/*lrp4* mutants as well as nerve-deprived pectoral fins revealed evidence for a previously unappreciated role for Agrin/Lrp4 signaling. While fins lacking Agrin or Lrp4 display reduced number of neural AChR clusters, the clusters that form are significantly enlarged in their median size, over 4-fold in *agrin* mutants (Fig. 5D). Strikingly, without Agrin or Lrp4, the giant AChR giant clusters can be partially suppressed by depleting *musk*. A developmental timecourse suggests these giant clusters are derived from MuSK-dependent prepatterned AChR clusters that expand over time. Combined our findings strongly support the idea that Agrin and Lrp4 play a dual role to both promote the formation of axon-induced AChR clusters and unexpectedly, to constrain Agrin-independent MuSK activity thereby restricting the growth and development of initial aneural clusters into synapse-associated AChR clusters. Future studies will be necessary to identify the downstream signaling events that allow Agrin/Lrp4 signaling to simultaneously potentiate and attenuate MuSK signaling in appendicular muscle at different regions on the same muscle fiber.

### Presynaptic axon patterning in appendicular muscle

Navigating growth cones respond to local extrinsic cues to determine where to form a synapse. This ‘stop signal’ requires MuSK signaling, as *musk* mutants in mouse and fish have an overgrown, defasciculated axon pattern (DeChiara et al., 1996; Zhang et al., 2004; Kim and Burden, 2008). In the zebrafish, this axon patterning role for MuSK is independent of AChR clustering, as *rapsyn* mutants that lack clustered AChRs have normal axon patterning (Zhang et al., 2004; Gribble et al., 2018) (Fig. S2). Thus, MuSK independently clusters AChRs postsynaptically and regulates axon patterning presynaptically, although the mechanism is poorly understood. Unexpectedly, we find that in the pectoral fin the presynaptic consequence of *agrin* or *lrp4* loss is for multiple axons to form mature synapses in apposition to postsynaptic AChR giant clusters, suggesting an overactive ‘stop signal’ in these regions. These giant cluster regions likely represent ‘islands’ of enhanced MuSK activity, as depletion of *musk* suppresses the formation of presynaptic swellings. Taken together, these results suggest that Lrp4 signaling, induced by the arrival of axonally released Agrin, may inhibit the MuSK-dependent ‘stop signal’ in the appendicular muscle of the pectoral fin. Such a mechanism could be a way to signal to new waves of navigating growth cones that this synaptic region is occupied.

Our data demonstrate that MuSK plays distinct roles to establish both the presynaptic axonal pattern and postsynaptic AChR clusters, but that both processes are constrained by Agrin/Lrp4 signaling within appendicular muscle. One attractive candidate for how MuSK might influence neuromuscular synapse development in an Agrin/Lrp4-dependent manner is through Wnt signaling. MuSK can bind Wnts through its extracellular cysteine-rich domain (CRD) (Jing et al., 2009; Strochlic et al., 2012; Zhang et al., 2012b). Indeed, in the zebrafish trunk, Wnt4a and Wnt11r binding through the MuSK CRD are required for AChR prepatterning and axon guidance (Jing et al., 2009; Gordon et al., 2012). While a functional role for Wnt:MuSK signaling to establish prepatterning in mouse is more controversial (Messéant et al., 2015; Remédio et al., 2016), in mammals Wnt proteins do regulate both AChR clustering (Strochlic et al., 2012; Zhang et al., 2012a) and axon guidance (reviewed in Zou, 2004). Interestingly, as the MuSK CRD domain can adopt two distinct conformations with differential abilities to bind Wnts (Stiegler et al., 2006), Guarino *et al*. speculate that Agrin/Lrp4 binding to MuSK can promote a conformational change to make MuSK unable to bind Wnts thereby shifting downstream MuSK signaling (Guarino et al., 2020). Therefore, it is tempting to speculate that in the pectoral fin prior to axon innervation, MuSK:Wnt signaling promotes both prepatterning of AChRs and local ECM cues to ‘catch’ navigating growth cones at future synaptic sites, perhaps through noncanonical Wnt signaling (Jing et al., 2009). Once the axon arrives, it releases Agrin, which binds Lrp4 on the muscle membrane and induces a conformational change in MuSK such that it can no longer bind Wnts. Thus, via competition through binding, Agrin/Lrp4 could locally constrain MuSK signaling. An outstanding question is why Agrin/Lrp4 regulate MuSK signaling differently between axial and appendicular muscle. As there are 23 Wnts in the zebrafish genome (Lu et al., 2011) with dynamic and differential expression throughout larval development, perhaps the relevant Wnt ligand is expressed in the pectoral fin but not in the trunk. Of course, there are many additional proteins that interact directly or indirectly with Agrin, Lrp4, or MuSK to influence synaptic development. Independently of the underlying mechanism, our results uncover a novel role for Agrin/Lrp4 signaling to constrain MuSK activity in appendicular muscle. The noncanonical nature of our findings reveal that there is diversity in the molecular pathways that mediate neuromuscular synapse development and validate the application of the larval zebrafish pectoral fin to study these processes beyond axial muscle.

## METHODS

### Zebrafish strains and animal care

Protocols and procedures involving zebrafish (*Danio rerio*) are in compliance with the University of Pennsylvania Institutional Animal Care and Use Committee regulations. All transgenic lines were maintained in the Tübigen or Tupfel long fin genetic background and raised as previously described (Mullins et al., 1994). The following transgenic lines were used: *Tg(mnx1:GFP)^ml2^* (Flanagan-Steet et al., 2005), *Tg(α-actin:Lrp4-GFP)^p159Tg^* (Gribble et al., 2018), *Tg(Xla.Tubb:DsRed)^zf148^* (Peri and Nüsslein-Volhard, 2008), *Gt(dmd-citrine)^ct90a^* (a kind gift from Dr. Sharon Amacher) (Ruf-Zamojski et al., 2015), and *Tg(α-actin:GFP)* (Higashijima et al., 1997). The following mutant strains were used: *agrin^p168^* (Gribble et al., 2018)*, lrp4^p184^* (Remédio et al., 2016), *lrp4^mi36^* (Saint-Amant et al., 2008), *musk^tbb72^* (Granato et al., 1996; Zhang et al., 2004), and *rapsyn*(*two^th26^*) (Ono et al., 2002). Homozygous mutants for these genes can be phenotyped at ~36 hpf as they all display motor defects when prodded with a probe. The *lrp4^p184^, two^th26^*, and *agrin^p168^* alleles were genotyped using Kompetitive Allele Specific PCR (KASP, LGC Biosearch Technologies). Animals were staged as previously published (Grandel and Schulte-Merker, 1998; Kimmel et al., 1995). As our experiments in larval zebrafish occur prior to sex determination, sex was not a biological variable (Kossack and Draper, 2019).

### Whole-mount immunohistochemistry and imaging

Zebrafish embryos or larvae immobilized with tricaine (MS-222) and then were fixed for 1 hour at room temperature in sweet fix (4% paraformaldehyde with 125mM sucrose in PBS) plus 0.1% Triton X-100 (Fisher, BP151). Animals were washed in phosphate buffer and incubated overnight at 4°C in primary chicken anti-GFP antibody (1:2000, Aves labs, GFP-1010) or mouse anti-Znp-1 (1:200, Developmental Studies Hybridoma Bank) in incubation buffer (2 mg/mL BSA, 0.5% Triton X-100, 1% NGS). After washing in phosphate buffer, animals were incubated overnight at 4°C in Alexa Fluor 488 donkey anti-chicken secondary antibody (1:1000, Jackson ImmunoResearch, 703-545-155), Alexa Fluor 594 goat anti-mouse secondary antibody (1:1000, Invitrogen, A-21125) and/or alpha-bungarotoxin Alexa Fluor 594 (1:500, Molecular Probes, B-13423) in incubation buffer. Animals were mounted in agarose in a glass-bottomed dish and imaged in 1.5 μm slices using a x40 or x63 water immersion lens on a Zeiss LSM880 confocal microscope using Zen software (Fig. 2C and 8) or a x40 water immersion lens on an ix81 Olympus spinning disk confocal microscope using Slidebook Software.

### Sparse neuronal labeling

A DNA vector encoding *mnx1:mKate* was injected as previously described (Gribble et al., 2018; Thermes et al., 2002) into one-cell-stage embryos. Embryos were screened at 1 dpf for larvae expressing mKate sparsely in the anterior spinal cord. At 5dpf, animals were mounted in agarose and imaged live at x40 on an Olympus spinning disk confocal if they had sparse mKate-expressing axons innervating the pectoral fin.

### Timelapse imaging

Embryos expressing both *α-actin:GFP* and *Xla.Tubb:DsRed* were anesthetized with tricaine and mounted in agarose around 35hpf. Animals were timelapsed using a 40x lense on an ix81 Olympus spinning disk confocal in a temperature chamber set to 28°C as previously described (Rosenberg et al., 2012). Stacks through the developing fin bud were captured in 1.5 μm slices with 30 minute intervals. Animals were imaged continuously for up to 3 days, with some adjustments to account for drifting and the pectoral fin moving out of frame.

### Motor neuron ablation

*mnx1:GFP* animals were mounted in agarose and motor neurons from spinal cord segments 3-5 were ablated using an Ablate! 532 nm attenuable pulse laser (Intelligent Imaging Innovations (3I), Denver, CO) beginning at 2 dpf, prior to axons innervating the pectoral fin bud. Neurons were considered ablated when there were no GFP+ cell bodies present in the ablated spinal cord region and axons showed signs of fragmentation. Neurons were re-ablated at 3 dpf to ensure any regenerated motor neurons in the spinal cord did not innervate the fin. Fins were visually inspected to confirm absence of GFP+ motor axon signal within the denervated region of the fin. Animals were fixed and stained with alpha-bungarotoxin at 5 dpf.

### Image processing and quantification

To simplify data visualization and quantification, signal from the abductor or adductor innervation of was manually separated from stacks through pectoral fins. Individual channel image stacks were opened in Fiji (Schindelin et al., 2012), background subtracted, channels were merged, and the image was changed to RGB. Stacks were visualized using the 3D viewer plugin and rotated to a top-down view so the separation between abductor and adductor innervation was distinct. Axon signal from the opposite innervation field and other fluorescent signal from the larval body wall was selected and filled, resulting in the corresponding region filled with black on the RGB stack. Any residual signal from the opposing innervation field or trunk was removed directly on the RGB stack. This resulted in signal specific to the abductor or adductor muscles, as specified. Stacks were converted to maximum projections. For quantification of α-Btx puncta, a custom CellProfiler (Lamprecht et al., 2007) pipeline was created to detect and measure the area of α-Btx punta per fin. Mutants that do not form distinct α-Btx puncta (*musk* and *rapsyn*) could not be quantified using CellProfiler pipelines. Fin images were excluded from analysis if the maximum projection did not include the whole fin, the fin was damaged or abnormally small, or if measurements were clear outliers from other fins of the same genotype in the dataset. All figures show only abductor innervation except for early developmental stages in figures 2 and 7, which are a maximum projection of the entire pectoral fin bud or where otherwise noted.

### Statistical analysis

Data were imported into Graphpad Prism for statistical analysis. Groups were compared using an unpaired t-test or one-way ANOVA with either Dunnett’s or Sidak’s multiple comparisons tests. For histogram of cluster sizes, the area of all AChR clusters measured in each genotype were pooled and binned into 5 μm^2^ bins, with any cluster over 20 μm^2^ included in the same bin. Distributions were compared using a Kruskal-Wallis test with Dunn’s multiple comparisons test. In figures where the control group is labeled as “siblings” we pooled wild type and heterozygous animals together as there was no statistical difference between these groups.

## ACKNOWLEDGEMENTS

The authors would like to thank members of the Granato lab, the UPenn zebrafish community, and the UPenn neurodevelopment group for helpful feedback on data and the manuscript. We would also like to thank Dr. Jonathan Raper for helpful discussions. Finally, we would like to thank the Penn Sanger sequencing core for technical support and the Penn Cell and Developmental Biology microscopy core.

## COMPETING INTERESTS

No competing interests declared.

## FUNDING

This work was supported by the National Institute of Neurological Disorders And Stroke of the National Institutes of Health under Award Numbers K01NS119496 (LW), F32 NS103219 (LW), R01NS097914 (MG) and RO1 EY024861 (MG).

## Notes

### Competing Interest Statement

The authors have declared no competing interest.

